# Inhibition of EGFR/ErbB does not protect against *C. difficile* toxin B

**DOI:** 10.1101/2024.05.13.594035

**Authors:** Uswah Siddiqi, Hannah M. Lunnemann, Kevin O. Childress, John A. Shupe, Stacey A. Rutherford, Melissa A. Farrow, M. Kay Washington, Robert J. Coffey, D. Borden Lacy, Nicholas O. Markham

**Affiliations:** Department of Medicine, Division of Gastroenterology, Hepatology, and Nutrition, Vanderbilt University Medical Center, Nashville, TN; Department of Pathology, Microbiology, and Immunology, Vanderbilt University Medical Center, Nashville, TN; Epithelial Biology Center, Vanderbilt University Medical Center, Nashville, TN; Department of Veterans Affairs, Tennessee Valley Healthcare System, Nashville, TN

## Abstract

*Clostridioides difficile* is a common cause of diarrhea and mortality, especially in immunosuppressed and hospitalized patients. *C. difficile* is a toxin-mediated disease, but the host cell receptors for *C. difficile* toxin B (TcdB) have only recently been revealed. Emerging data suggest TcdB interacts with receptor tyrosine kinases during infection. In particular, TcdB can elicit Epidermal Growth Factor Receptor (EGFR) transactivation in human colonic epithelial cells. The mechanisms for this function are not well understood, and the involvement of other receptors in the EGFR family of Erythroblastic Leukemia Viral Oncogene Homolog (ErbB) receptors remains unclear. Furthermore, in an siRNA-knockdown screen for protective genes involved with TcdB toxin pathogenesis, we show ErbB2 and ErbB3 loss resulted in increased cell viability. We hypothesize TcdB induces the transactivation of EGFR and/or ErbB receptors as a component of its cell-killing mechanism. Here, we show in vivo intrarectal instillation of TcdB in mice leads to phosphorylation of ErbB2 and ErbB3. However, immunohistochemical staining for phosphorylated ErbB2 and ErbB3 indicated no discernible difference between control and TcdB-treated mice for epithelial phospho-ErbB2 and phospho-ErbB3. Human colon cancer cell lines (HT29, Caco-2) exposed to TcdB were not protected by pre-treatment with lapatinib, an EGFR/ErbB2 inhibitor. Similarly, lapatinib pre-treatment failed to protect normal human colonoids from TcdB-induced cell death. Neutralizing antibodies against mouse EGFR failed to protect mice from TcdB intrarectal instillation as measured by edema, inflammatory infiltration, and epithelial injury. Our findings suggest TcdB-induced colonocyte cell death does not require EGFR/ErbB receptor tyrosine kinase activation.

## Introduction

*Clostridioides difficile* (*C. difficile)* is a toxin-producing, Gram-positive bacillus and is a major cause of healthcare-associated diarrhea and mortality [1]. The virulence factors causing most epithelial damage and inflammation are two large exotoxins, toxin A (TcdA) and toxin B (TcdB) [2]. The contribution of a third toxin, transferase toxin (CDT), is not as well-understood [3]. TcdA and TcdB have four functional domains: the glucosyltransferase domain (GTD), the auto-protease domain (APD), the delivery domain, and the combined repetitive oligopeptides (CROPS) domain [4,5]. Both toxins interact with multiple cell surface receptors (discussed below), and they are internalized into endosomes [6]. Acidification of the endosome triggers a conformational change in the toxin delivery domain, leading to pore formation and the translocation of the APD and GTD into the cytosol [7]. The auto-proteolytic cleavage leads to the release of the GTD so it can inactivate small RhoGTPases: RHO, RAC, and CDC42 via mono-glucosylation. These proteins are guanosine triphosphatases that regulate cytoskeletal dynamics, cell adhesion, and signal transduction [8]. Their inhibition results in cell rounding and apoptosis of the intoxicated host cell [9].

Some evidence has suggested TcdA is essential for *C. difficile* infection (CDI) in humans [10], but more recent data from patients infected with strains lacking TcdA and CDT demonstrate TcdB alone can cause epithelial cell damage, inflammation, and a full range of clinical symptom severity [11]. Interestingly, high concentrations (> 0.1 nM) of TcdB can induce glucosyltransferase-independent necrosis through over-production of reactive oxygen species [12]. Moreover, bezlotoxumab, a neutralizing antibody against TcdB, is superior to actoxumab, a neutralizing antibody against TcdA, for reducing the rate of recurrent CDI [13]. In pre-clinical models, isogenic knockout strains of *C. difficile* have shown TcdB is required for inducing wild-type levels of tissue damage and mortality [14–16].

TcdB receptors include chondroitin sulfate proteoglycan 4 (CSPG4), Nectin 3 (PVRL3), frizzled proteins (FZDs), and tissue factor pathway inhibitor (TFPI) [17]. FZDs are essential receptors for WNT ligands that promote proliferation and self-renewal of colonic epithelial cells [18]. The interaction between TcdB and FZDs, particularly FZD1, 2, and 7, may block WNT signaling and contribute to cell death [19,20].

TcdB has been reported to induce EGFR transactivation as a part of the cell death mechanism in CDI [21]. Specifically, in the non-transformed human colonic epithelial cell line, NCM460, TcdB promotes TGFα-dependent phosphorylation of EGFR, and subsequent activation of the ERK/MAP kinase cascade leads to IL-8 cytokine production. However, these data cannot rule out the contribution of other ErbB receptors or address whether receptor transactivation contributes significantly to cell death. EGFR and ErbB2 are essential for cell survival via anti-apoptotic signals in mouse colonocytes in the setting of acute inflammation, but their role in the setting of TcdB-induced inhibition of small RhoGTPases during CDI is unclear. CDC42-deficient intestinal organoids undergo rapid apoptosis because CDC42 engaging with EGFR is required for EGF-stimulated receptor-mediated endocytosis. Interestingly, treating breast cancer cells in vitro with TcdB results in altered ErbB2 expression patterns leading to decreased tumor burden [22].

Receptor tyrosine kinase inhibitors (TKIs) have found wide application in treating solid tumors and hematological malignancies, effectively blocking signaling pathways that drive tumor growth and spread [23]. Emerging clinical evidence suggests TKIs could potentially have a protective role against CDI. In a study assessing the effect of anti-EGFR TKIs on CDI, lung cancer patients with diarrhea due to *C. difficile* had a longer interval between TKI initiation and diarrhea (median period: 75 days, range: 25 to 376 days) compared with patients who had diarrhea due to other causes (median period: 7 days, range: 0 to 49 days) [24]. The incidence of CDI in this study was 2.2%, which is notably lower compared to another study of cancer patients receiving immune checkpoint inhibitor immunotherapy where the incidence of CDI was 9.7% [25]. One possible explanation for these results is that blocking EGFR and/or ErbB receptor signaling is protective against CDI. We hypothesize transactivation of EGFR and ErbB receptors is increased by TcdB and is a component of its cell-killing mechanism during CDI.

Herein, we demonstrate that siRNA-knockdown of ErbB2 or ErbB3 protects against TcdB-mediated cell death, and ErbB2 and ErbB3 are phosphorylated selectively by TcdB in the mouse colon. However, phospho-ErbB2 and phospho-ErbB3 staining in mouse colonic epithelium by immunohistochemistry show no significant differences in abundance or localization between TcdB- and vehicle-instilled mice. In vitro, HT-29 cells did not show co-localization of phosphor-EGFR and TcdB, nor were HT-29 or Caco-2 cells protected against TcdB when pre-treated with lapatinib, a potent EGFR/ErbB2 inhibitor. To eliminate the possible confounding factors associated with using cancer cell lines, we performed TcdB-intoxication experiments on normal human colon organoids (colonoids). These colonoids were not protected from TcdB by either lapatinib or a more general tyrosine kinase inhibitor dasatinib. Finally, we expanded our investigations to include neutralizing antibodies against EGFR and found that this form of inhibition did not protect the mouse colon or human colonoids from TcdB.

## Methods

### siRNA knockdown screen

HeLa cells were seeded into 96-well plates with a human siRNA knockdown library (Dharmacon) and transfection reagent per manufacturer’s protocol. Cells were incubated with TcdB (100 ng/mL) at 37 °C, and cell viability was measured with CellTiter-Glo luciferase assay. Results are expressed as viability relative to non-targeting control siRNA. TcdB was prepared recombinantly as described [26].

### Phospho-specific receptor activation assay

Wild-type C567Bl/6 female mice aged 8-10 weeks (Jackson Labs) were acclimatized to our AAALAC-accredited animal facility for 2 weeks. All animal experiments were performed humanely under the IACUC-approved protocol #V2100012. Mice were anesthetized and instilled intrarectally with purified, recombinant TcdB as previously described [27]. After 4 h, mice were euthanized and whole colon tissue was harvested and flash frozen. Only the most distal 4 cm of colon were used. Whole tissue lysates were prepared as described previously [28]. Lysates of 3 mouse colons from each group were pooled together and used as a substrate for the mouse Proteome Profiler Array (R&D Systems) following the manufacturer’s protocol.

### Immunohistochemistry

Following intrarectal instillations as detailed above, whole mouse colons were harvested, washed, formalin-fixed, and paraffin-embedded. Tissue blocks were sectioned and prepared for immunohistochemistry (Agilent Technologies) as previously described [27]. Antibodies used were: total-ErbB2 clone D8F12, phospho-ErbB2 clone 6B12, total ErbB3 clone D22C5, and phospho-ErbB3 clone 21D3 (Cell Signaling Technologies).

### Immunofluorescence

HT-29 cells (kindly gifted by the Coffey Lab) were seeded into 12-well plates containing sterile glass coverslips. After reaching 90-100% confluence, TcdB was added to the cells in fresh DMEM media (Gibco). Recombinant human EGF (R&D Systems) was applied similarly as a positive control. Cells were washed and fixed to the coverslips at the indicated times with 10% neutral buffered formalin (Sigma). Coverslips were permeabilized, blocked, and immunostained using previously published methods [29]. The phospho-EGFR antibody (clone 53A5, Cell Signaling) and anti-TcdB sheep polyclonal antibody (R&D Systems) were used with species-specific secondary antibodies (Fisher Scientific).

### Cell viability assay

HT-29 and Caco2 human adenocarcinoma cells were seeded in 96-well plates in triplicate and grown to near confluency. Cells were washed three times in Hank’s balanced salt solution, then incubated with lapatinib at the indicated concentrations at 37 °C. TcdB was added at the indicated concentrations after 1 hour and incubated at 37 °C for 20 h. Viability was measured with CellTiter-Glo (Promega) and normalized to untreated controls.

### Normal human colonoid viability assays

Normal human colon tissue was obtained from normal-adjacent surgical resection tissue through the Cooperative Human Tissue Network under the IRB-exempt protocol for Dr. Lacy. Epithelium was carefully dissected from the submucosa using sterile technique. Colonoids were derived as previously described for mouse enteroids [30] with the exception that chelation was performed with 50 mM EDTA/EGTA in PBS and 1 h incubation time. Colonoids were passaged no more than 7 times prior to these experiments. Colonoids were seeded into Matrigel (Corning) domes at a density of 100 cells/μL and cultured for 7-10 days prior to the experiment. Cells were pretreated with small molecule inhibitors lapatinib or dasatinib (Tocris) dissolved in 1% dimethyl sulfoxide and 99% Dubelco’s Modified Eagle Medium (DMEM). Experiments using EGFR neutralization were performed with C225 monoclonal antibody (kindly provided by the Coffey Lab) diluted in DMEM. TcdB was diluted in DMEM for addition to the colonoids after 1 h pretreatment. At the indicated times, colonoids were imaged with phase-contrast light microscopy (Keyence BZ-X800) for counting or lysed for viability assays using CellTiter-Glo 3D (Promega) and a luminometer (Bio-Tek Synergy HTX).

### EGFR neutralization in vivo

P1X/P2X antibodies (Merrimack Pharmaceuticals) were diluted in PBS to a concentration of 2.5 mg/mL and injected intraperitoneally at a dose of 25 mg/kg 3 times over 5 days to establish a steady-state in vivo concentration. Control mice were injected with an equal volume of sterile phosphate-buffered saline. On the 6^th^ day, mice were instilled with 5 or 50 μg TcdB as previously published and euthanized after 4 or 18 h [27]. Mouse colons were prepared as formalin-fixed, paraffin-embedded tissue as described above and sectioned for H&E staining using standard techniques. Dr. Washington reviewed and scored the tissue blindly using a previously published scoring rubric[31]. Colon tissue sections were prepared for immunofluorescence staining as previously described [27] using goat anti-human-AF647 and rabbit anti-GFP-AF488 conjugated antibodies (ThermoFisher, Invitrogen).

## Results

To identify host factors contributing to TcdB-mediated cell killing, we analyzed results from an in vitro siRNA-knockdown screen in HeLa cells treated with TcdB. Compared with non-targeting controls, the siRNAs against ErbB2 and ErbB3 led to increased viability (Figure 1A). The siRNAs against ErbB4 and the EGF-domain-containing transmembrane protein CD97 did not have any effect on viability. Rac and 6V0C are known components of the TcdB-pathogenesis mechanism and serve as positive controls.

**Figure 1.**
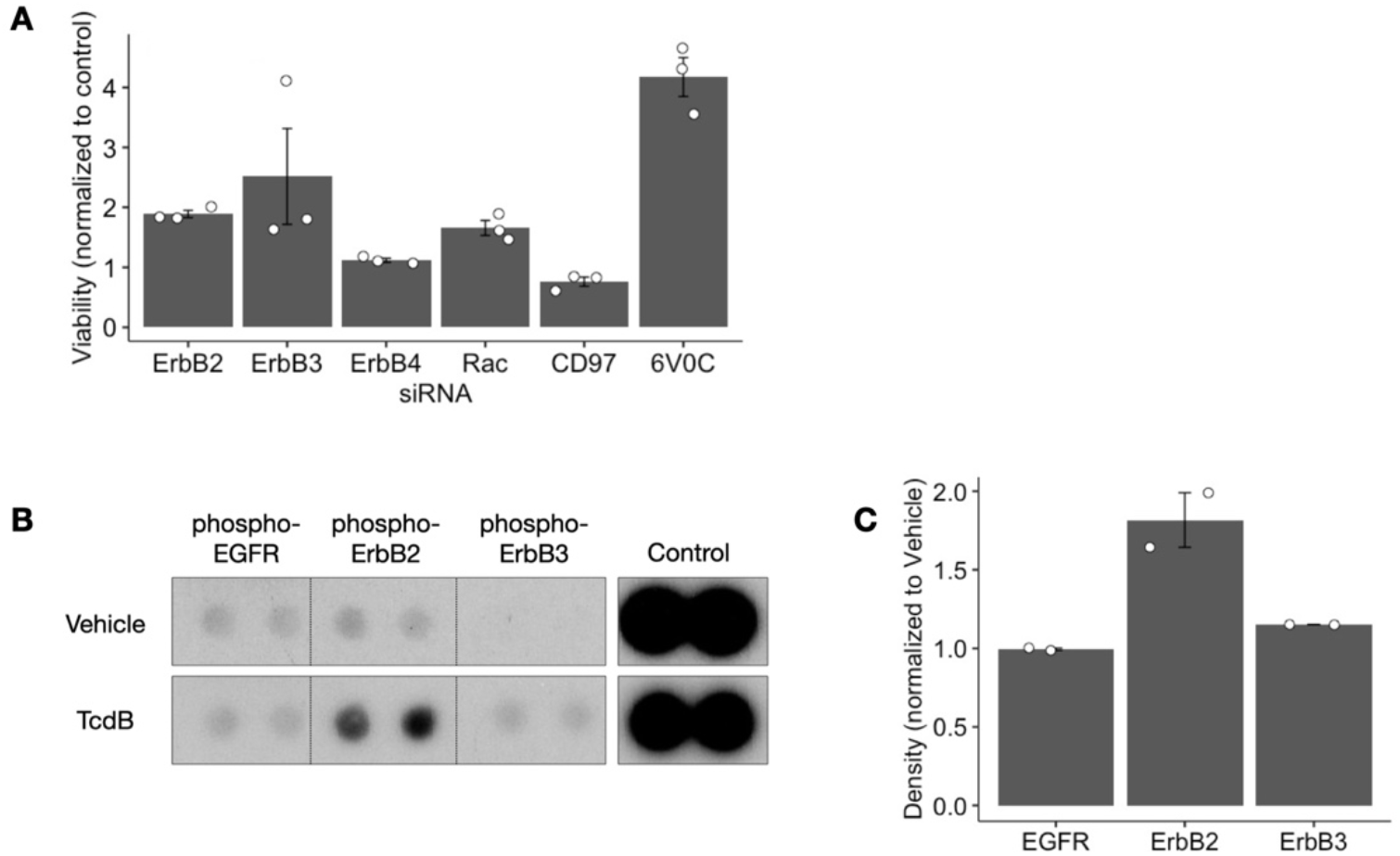
ErbB2 and ErbB3 are identified as potential contributors to TcdB pathogenesis from 2 screening experiments. A) siRNA-knockdown of ErbB2 and ErbB3 increased the relative viability of HeLa cells exposed to 100 ng/mL TcdB. B) Dot blot from receptor tyrosine kinase phospho-proteome array shows ErbB2 and ErbB3 are phosphorylated in pooled whole tissue lysates of 3 mouse colons 4 h after 50 μg TcdB intrarectal instillation. C) Densitometry of the dot blot in (B) shows increased ErbB2 and ErbB3 phosphorylation but not EGFR phosphorylation.

In a parallel screen using intrarectal instillation of TcdB in the mouse colon, we measured the phosphorylation of receptor tyrosine kinases. Purified recombinant TcdB or vehicle control were intrarectally instilled into mouse colons, and whole distal colon tissue was harvested after 4 h. Colon samples were pooled based on TcdB instillation or vehicle control (n=3 mice/group). ErbB2 and ErbB3 were the only phosphorylated receptors in the mouse colon in TcdB-exposed versus vehicle controls (Figure 1B).

To determine the location and cell type-specific expression of phospho-ErbB2 and phospho-ErbB3 and to validate the screening results, we instilled wild-type C57Bl/6 female mice with purified recombinant TcdB (50 μg) or vehicle control. We compared the amount of immunohistochemistry staining for phospho-ErbB2 and phospho-ErbB3 as well as total ErbB2 and total ErbB3 (Figure 2A-B). From 6-8 mice per group, we did not observe any differences in staining between TcdB- and vehicle-instilled colons among any of the ErbB antigens.

**Figure 2.**
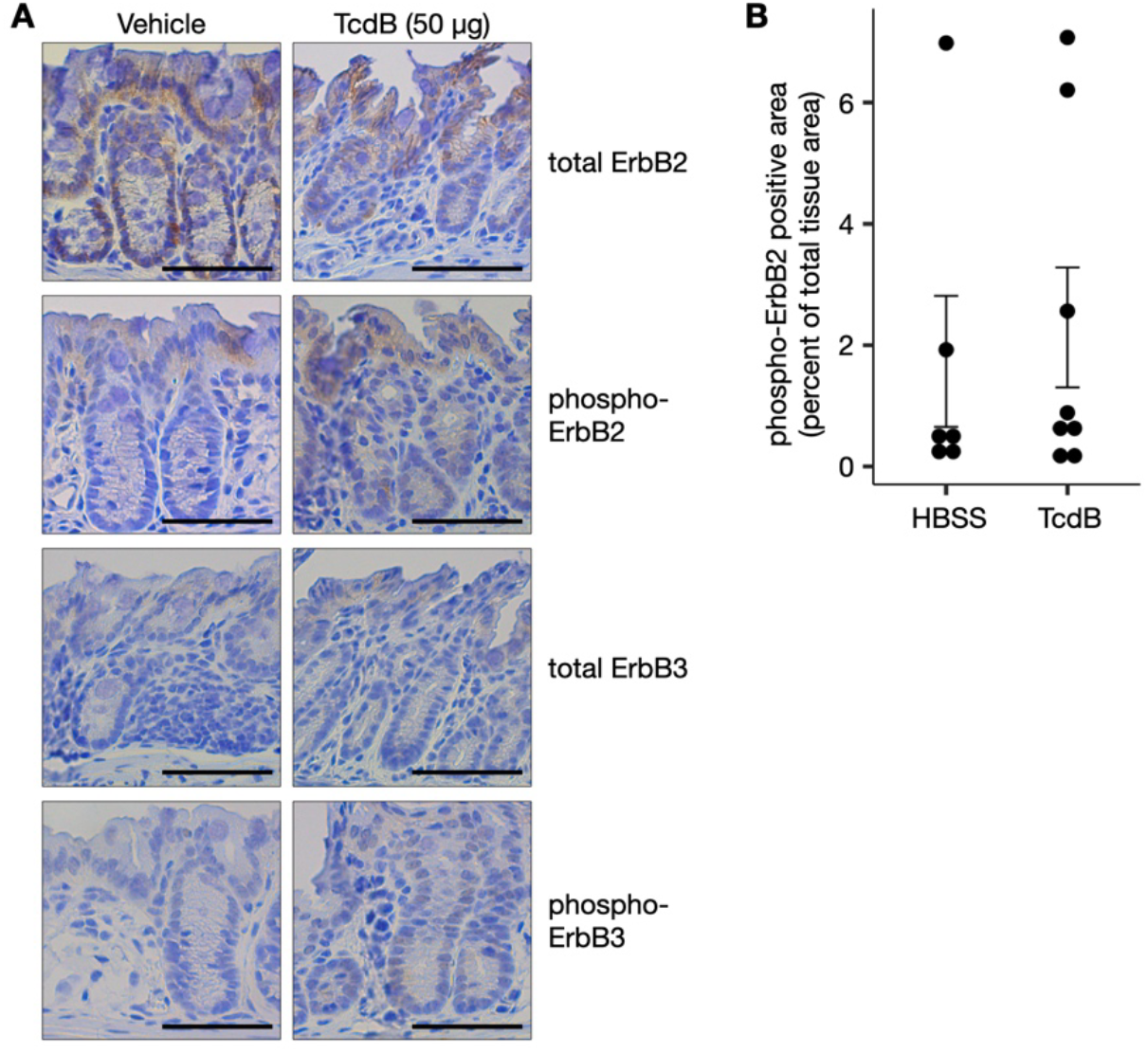
ErbB2 and ErbB3 phosphorylation in TcdB-instilled mouse colon. A) Immunohistochemistry staining shows expression of 4 antigens (total ErbB2, phospho-ErbB2, total ErbB3, and phospho-ErbB3) in distal mouse colon 4 h after instillation with either purified recombinant TcdB or vehicle control; scale bar = 50 μm. B) Quantification of the immunohistochemistry in (A) shows no statistical different using the Mann-Whitney U test.

Next, we wanted to see if TcdB induces phosphorylation of EGFR in HT-29 human colon cancer cells. Using a specific antibody against EGFR phospho-tyrosine 1173 (pY1173), we performed immunofluorescence at 15-120 minutes after exposure to 10 nM of TcdB. At 30 minutes, there is increased staining for pY1173 diffusely but not precisely co-localizing with the highest abundance of TcdB at specific cell-cell junctions (Figure 3A). Detection of pY1173 is not seen at other time points. To determine if inhibition of EGFR and ErbB2 transactivation protects HT-29 or Caco2 cells, we performed viability assays over a range of TcdB concentrations after pre-treatment with lapatinib, a specific and potent inhibitor of the EGFR and ErbB2 intracellular ATP-binding site [32]. Under these conditions, lapatinib does not have a significant effect on the cell viability of either HT-29 or Caco2 cells exposed to TcdB (Figure 3B-C).

**Figure 3.**
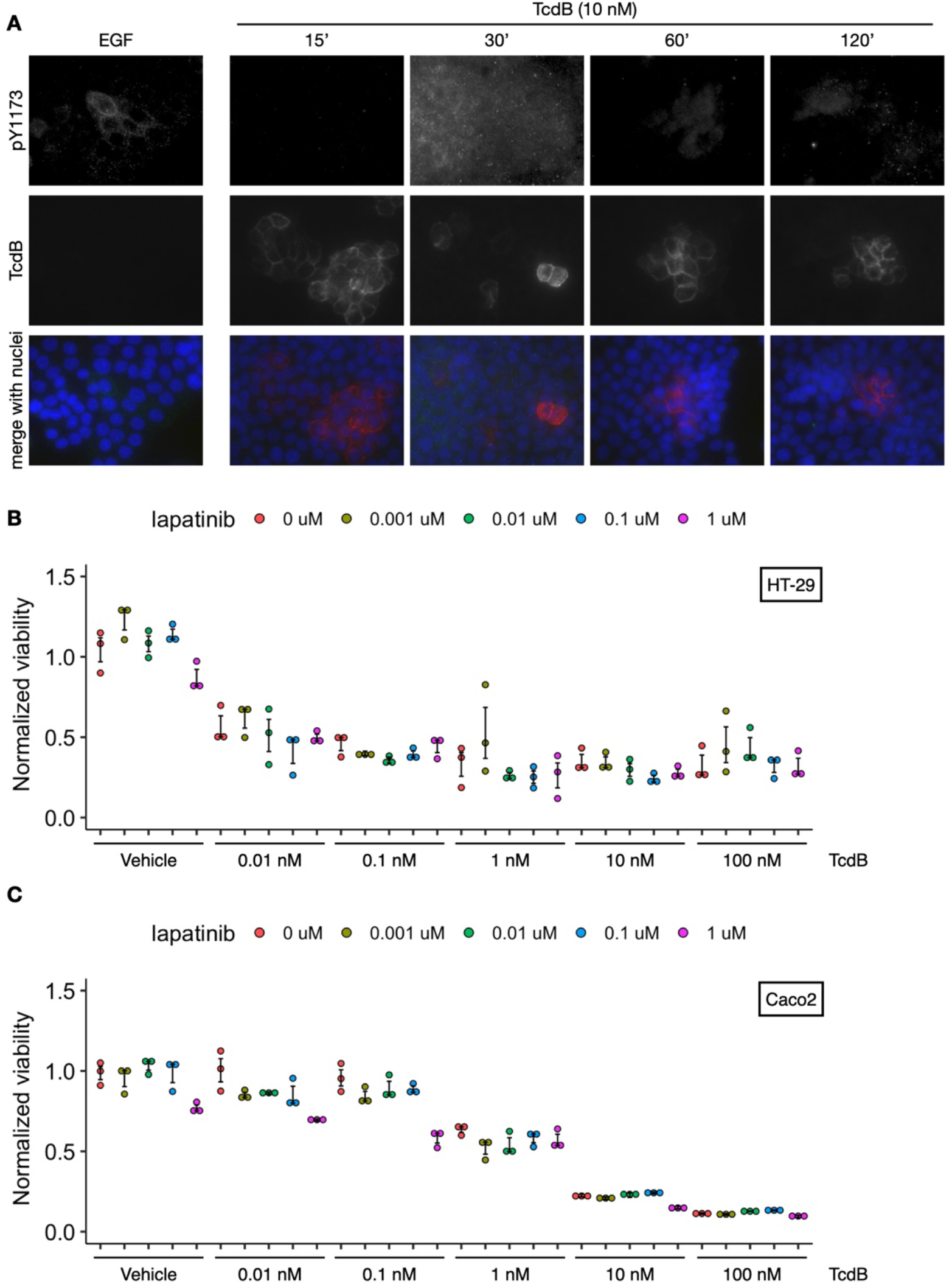
HT-29 and Caco2 human adenocarcinoma cells are not protected from TcdB by lapatinib pretreatment. A) Immunofluorescent staining of HT-29 cells treated with TcdB (10 nM) for 15-120 minutes shows phosphorylation of EGFR at tyrosine Y1173 and localization of TcdB. Epidermal growth factor (EGF) is used as a positive control. B, C) Cell viability assays for HT-29 and Caco2 cells using CellTiter-Glo show lapatinib does not protect cells from TcdB. Within any given TcdB concentration, there was no significant difference (adjusted *p*-value < 0.05) between lapatinib and vehicle or between different lapatinib concentrations as measured by one-way Kruskal-Wallis with Dunn’s post-hoc test for multiple comparisons and Bonferroni correction.

Adenocarcinoma cell lines like HT-29 and Caco2 grown on 2-dimensional plastic may have changes in receptor tyrosine kinase expression or dependency that could interfere with modeling the effect of TcdB in normal, non-cancerous cells [33]. Therefore, we derived organoids from normal human colon biopsies taken during routine screening colonoscopy. These normal human colonoids were grown as 3-dimensional spheres in Matrigel extracellular matrix and pretreated with lapatinib or dimethyl sulfoxide control. The same organoids were imaged iteratively over 22 h and viable organoids were counted. Decreased cell viability was apparent in TcdB-exposed colonoids compared to vehicle control (Figure 4A). Lapatinib pre-treatment did not improve cell viability compared with dimethyl sulfoxide (DMSO) controls (Figure 4A-B). Next, we tested lower concentrations of TcdB (10-20 pM) in normal human colonoids pretreated with lapatinib (4 μM) for 1 h. TcdB alone significantly decreased relative viability compared with DMSO controls (Figure 4C). However, cells pretreated with lapatinib were not protected against TcdB-mediated cell death. Ligand-independent phosphorylation of EGFR can occur through activation of SRC kinase [34]. To determine if this mechanism contributes to TcdB pathogenesis, we pretreated cells with dasatinib, a small molecule inhibitor of multiple tyrosine kinases, including SRC. Similarly to lapatinib, pretreatment with dasatinib did not improve the relative viability of TcdB-exposed normal human colonoids (Figure 4D). In fact, normal human colonoids given dasatinib alone had reduced viability compared to DMSO control.

**Figure 4.**
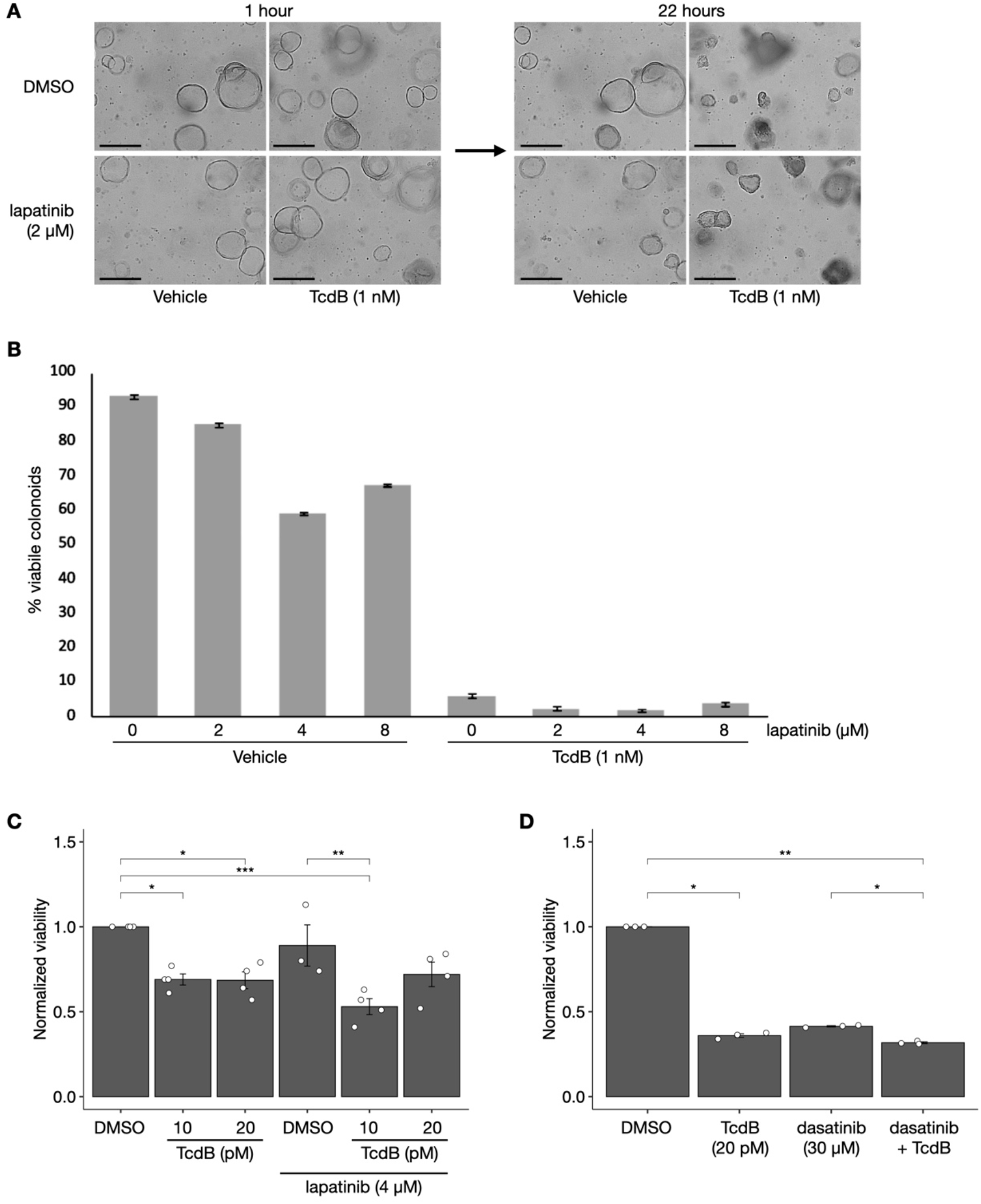
Lapatinib pretreatment does not protect normal human colonoids from TcdB-mediated cell death. A) Phase contrast light microscopy images show normal human colonoids are killed by TcdB (1 nM), but they are not protected by lapatinib (2 μM) given 1 h before TcdB; scale bar = 50 μm. B) Quantification of the colonoid experiment in (A) using a range of lapatinib concentrations. C) CellTiter-Glo viability assays with normal human colonoids, lapatinib pretreatment, and lower concentrations of TcdB (10-20 pM). D) CellTiter-Glo viability assay using dasatinib (30 μM) pretreatment, which does not protect colonoids from 20 pM TcdB.

To determine if the EGFR tyrosine kinase receptor participates in TcdB pathology in vivo, we performed in vivo mouse experiments using a pair of humanized neutralizing antibodies (P1X/P2X) against the mouse EGFR extracellular domain [35]. The recipient mice contained an Emerald-GFP fused to the C-terminus of the EGFR gene using CRISPR-Cas9 gene editing [36]. These mice were essential for this experiment, because reagents for detecting mouse EGFR are not specific. Animals were given intraperitoneal injections of P1X/P2X humanized antibodies for 5 days to obtain a steady-state tissue concentration. Then, mice were given intrarectal instillations of purified recombinant TcdB. Mouse colons were harvested 4 h after instillation and stained with H&E for blinded scoring by a gastrointestinal pathologist. No significant differences were observed between TcdB- and Vehicle-instilled mice with respect to edema, inflammatory infiltration, or epithelial injury (Figure 5A). The P1X/P2X pair of neutralizing antibodies appear to have been effective because we observed them in the colonic tissue by immunofluorescence (Figure 5B), and the EGFR-GFP fusion protein had reduced expression (Figure 5C).

**Figure 5.**
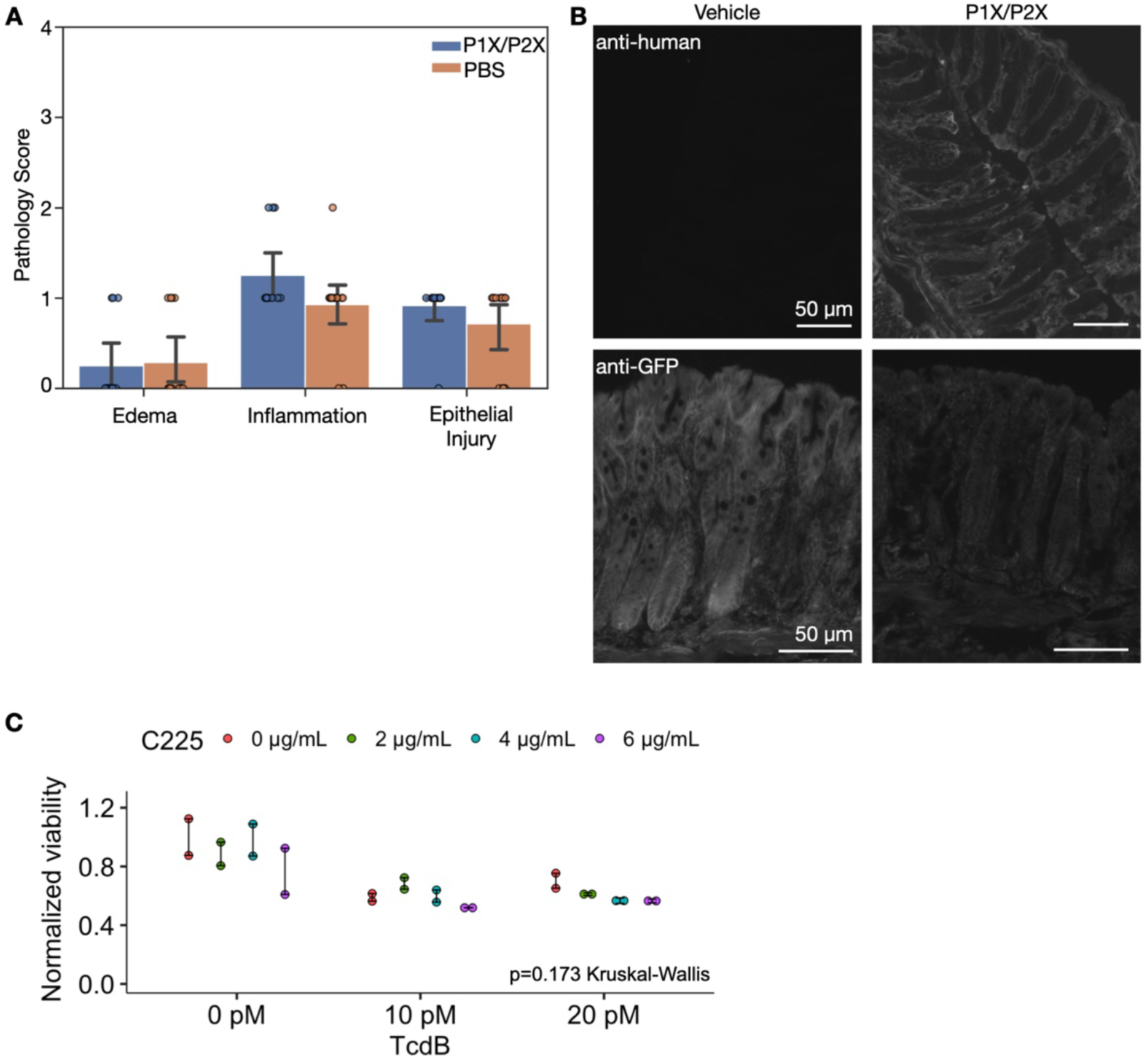
Neutralizing antibodies against mouse EGFR fail to protect the colon from TcdB-mediated damage. A) Blinded histopathological scoring of mouse colon H&E tissue from P1X/P2X-injected EGFR-EmeraldGFP mice compared to Vehicle controls. This bar plot represents the mean score with error bars showing 95% confidence intervals. *p*-values were calculated using the Mann-Whitney U test; Edema: 0.87, Inflammation: 0.10, Epithelial Injury: 0.21. B) Immunofluorescent staining of mouse colon tissue shows accumulation of the humanized, neutralizing antibodies (top row) in P1X/P2X-treated mice and efficacy of the neutralization as evident by reduction in EGFR-EmeraldGFP signal. C) TcdB induces human colonoid death at 10-20 pM over 24 hrs, but C225 neutralizing antibody against human EGFR does not protect colonoids; 2 independent wells, ∼200 organoids/well. Kruskal-Wallis test was used to identify a statistical difference between normalized viability as a function of C225 treatment, but the null hypothesis was confirmed (*p* = 0.173).

We performed a similar EGFR-neutralizing experiment in normal human colonoids to determine if there is a species-specific role for EGFR in TcdB-mediated cell death. In this experiment, we pretreated colonoids with the C225 antibody that neutralizes human EGFR similarly to the anti-cancer drug cetuximab, which inhibits ligand binding and subsequent receptor dimerization [37]. C225 was added to serum-starved normal human colonoids 1 h before TcdB. Cell-permeable Hoechst 33342 and propidium iodide (live cells are impermeable to propidium iodide) were then added to all colonoids 24 h after TcdB. To assess colonoid viability, fluorescence microscopy measured the intensity of propidium iodide and Hoechst. The C225 neutralizing antibody did not improve colonoid viability, which was calculated as the inverse of the propidium iodide-Hoechst ratio. Taken together, we ultimately found that specifically blocking EGFR and/or ErbB2 in multiple models does not alter the effect of TcdB on colonocyte death or injury.

## Discussion

The appearance of ErbB2 and ErbB3 as protective factors against TcdB-mediated cell death was initially surprising given the critical role of these receptors for epithelial restitution in intestinal damage and colitis [38,39]. These receptors are largely considered to promote cell growth, but their transactivation can lead to rapid cell death in the absence of the RhoGTPase CDC42 [40]. This led us to hypothesize that the EGFR family of tyrosine kinase receptors might activate apoptosis in the setting of TcdB glucosyltransferase-mediated inhibition of RhoGTPases. Previous studies have shown TcdB-induced phosphorylation of EGFR in human cell lines, but the reagents available at that time were not specific to EGFR and might have cross-reacted with other family members, including ErbB2 and ErbB3 [21].

Our initial screening experiments supported our hypothesis. Phospho-proteome array data suggested there might be a role for ErbB2 and ErbB3 in TcdB-induced mouse colitis, because they were specifically phosphorylated in whole tissue lysates (Figure 1B). Looking at the same mouse colon tissue with immunohistochemistry did not show any significant differences for ErbB2 or ErbB3 total or phosphorylated protein in the epithelium (Figure 2A-B). Perhaps this contradiction can be explained by increased ErbB2 and ErbB3 phosphorylation in stromal cells or lymphoid aggregates form the whole tissue lysates used in the phospho-proteome array that were not seen in the fixed tissue sections used for immunohistochemistry.

While TcdB appeared to increase the phospho-EGFR signal in HT-29 cells by immunofluorescence (Figure 3A), it was not seen in the same location at the cell membrane as TcdB. There was no protection against TcdB afforded by the pre-treatment of HT-29 or Caco-2 cells with lapatinib, a specific EGFR/ErbB2 inhibitor (Figure 3B-C). The detection of transactivation in these cell lines was not as obvious as reported phosphorylation in NCM460 cells [21]. The baseline expression of receptors and the ligands present in the fetal bovine serum may have been very different. Therefore, it was important for us to test different cell lines, including normal human colonoids, and different concentrations of TcdB with different tyrosine kinase inhibitors (Figures 4-5). Ultimately, we were unable to observe any functional effect of EGFR/ErbB inhibition during TcdB-mediated cell killing.

In summary, we undertook a series of experiments to determine the influence of EGFR/ErbB transactivation on TcdB toxin pathogenesis. The inhibition of EGFR and ErbB2/3 did not have a detectable affect on viability in our cell lines, organoids, or mouse model. While these receptors may influence TcdB in some circumstances, we have not found a robust effect.

## Acknowledgments

We wish to acknowledge the kind gift of C225 antibody from the laboratory of Robert J. Coffey, Jr, MD. Core Services performed through Vanderbilt University Medical Center’s Digestive Disease Research Center supported by NIH grant P30DK058404. Funding to N.O.M. and D.B.L. was provided by the U.S. Department of Veterans Affairs via grants BX005699 and BX002943, respectively. The contents do not represent the views of the United States Department of Veterans Affairs or the United States Government.

